# Disentangling the Innate Immune Responses of Intestinal Epithelial Cells and Lamina Propria Cells to *Salmonella* Typhimurium Infection in Chickens

**DOI:** 10.1101/2023.07.14.548977

**Authors:** Kate Sutton, Tessa Nash, Samantha Sives, Dominika Borowska, Jordan Mitchell, Prerna Vohra, Mark P. Stevens, Lonneke Vervelde

## Abstract

*Salmonella enterica* serovar Typhimurium (STm) is a major foodborne pathogen and poultry are a key reservoir of human infections. To understand the host responses to early stages of *Salmonella* infection in poultry, we infected 2D and 3D enteroids, the latter of which contains leukocytes, neurons, and mesenchymal cells that are characteristic of the lamina propria. We infected these enteroids with wild-type (WT STm), a non-invasive mutant lacking the *prgH* gene (*ΔprgH* STm), or treated them with STm lipopolysaccharide (LPS) and analysed the expression of innate immune related genes by qPCR at 4 and 8 h. The localisation of ZO-1 expression was disrupted in WT STm infected enteroids but not *ΔprgH* STm or LPS treated enteroids, suggesting a loss of barrier integrity. The innate immune response to LPS was more pronounced in 2D enteroids compared to 3D enteroids and by 8 hpi, the response in 3D enteroids was almost negligible. However, when STm adhered to or invaded the enteroids, both 2D and 3D enteroids exhibited an upregulation of inflammatory responses. The presence of lamina propria cells in 3D enteroids resulted in the unique expression of genes associated with immune functions involved in regulating inflammation. Moreover, 2D and 3D enteroids showed temporal differences in response to bacterial invasion or adherence. At 8 hpi, innate responses in 3D but not 2D enteroids continued to increase after infection with WT STm, whereas the responses to the non-invasive strain decreased at 8 hpi in both 2D and 3D enteroids. In conclusion, STm infection of chicken enteroids recapitulated several observations from *in vivo* studies of *Salmonella*-infected chickens, including altered epithelial barrier integrity based on ZO-1 expression and inflammatory responses. Our findings provide evidence that *Salmonella*-infected enteroids serve as effective models for investigating host-pathogen interactions and exploring the molecular mechanisms of microbial virulence although the 3D model mimics the host more accurately due to the presence of a lamina propria.

## 1. Introduction

*Salmonella enterica* are Gram-negative rod-shaped facultative anaerobic bacteria that are comprised of over 2,600 antigenically distinct serovars. *Salmonella enterica* serovar Typhimurium (STm), typically has a broad host range and transmits via contaminated food or water, causing severe gastroenteritis. The consumption of poultry meat and eggs contaminated with STm is a significant contributor to human infections. Intestinal inflammation that characterises *Salmonella* gastrointestinal infection is caused by the infection of effector proteins into host cells by a Type 3 secretion system (T3SS-1) encoded by *Salmonella* pathogenicity island 1 (SPI-1) (Mills et al., 1995). Effector proteins delivered by T3SS-1 promote bacterial invasion by orchestrating rearrangements in the subcortical actin cytoskeleton and activate inflammatory responses (Raffatellu et al., 2005; Boyle et al., 2006). In mammals, mutations in T3SS-1 genes, such as *prgH*, reduce the ability of STm to colonise the intestine and induce inflammatory and secretory responses (Klein et al., 2000). T3SS-1 contributes to colonisation of the avian intestine by STm (Chaudhuri et al., 2013). However, inflammation is less pronounced than in mammals, with STm typically being carried asymptomatically in chickens over one week old and shed persistently in the faeces (Raffatellu et al., 2005). Neonatal chicks are highly susceptible to STm infection, which causes systemic infection and death (Barrow et al., 1987; Withanage et al., 2005). Although adults are less susceptible, STm can colonise the gastrointestinal tract without an associated clinical disease.

*In vivo* studies have provided considerable knowledge about the nature and consequences of mucosal immune responses to STm in the chicken intestine (Withanage et al., 2004; Iqbal et al., 2005; Withanage et al., 2005; Fasina et al., 2008; Bescucci et al., 2022). An *in vitro* analysis of *Salmonella -* host interactions and the contribution of epithelial and lamina propria cells have been lacking due to no availability of an enterocyte cell line or a defined reproducible and robust intestinal culture system in chickens. Three-dimensional (3D) intestinal organoids, when derived from primary tissue are known as enteroids, closely mimic the morphology and physiology of the intestine, and are emerging as *in vitro* models to study host-pathogen interactions. Intestinal enteroids grown in an extracellular matrix consist of a central lumen lined by a single layer of highly polarized epithelial cells with their basolateral surface in contact with the extracellular matrix scaffold (Sato et al., 2009). In contrast to cell lines, enteroids recapitulate all major differentiated epithelial cell lineages, including enterocytes, goblet cells, enteroendocrine cells, Paneth cells, and tuft cells. Zhang et al. (2014) were the first to analyse STm infection in murine enteroids demonstrating epithelial cell invasion, disruption of tight junctions and NFκB related pro-inflammatory responses. Human, bovine and porcine enteroids have since been reported to be susceptible to STm (Zhang et al., 2014; Forbester et al., 2015; Derricott et al., 2019). However, the fully enclosed lumen of mammalian enteroids poses a challenge to deliver the pathogens to the epithelial surface. Recently, apical-out enteroids derived from basal-out human, porcine, bovine and ovine enteroids have been developed (Co et al., 2021; Smith et al., 2021; Blake et al., 2022; Joo et al., 2022). A study has shown that human apical-out enteroids recapitulate specific morphological hallmarks of STm infection in humans including epithelial barrier disruption and cytoskeletal reorganisation (Co et al., 2021).

Avian floating 3D enteroids naturally develop in an advantageous apical-out conformation with apical microvilli facing the media and an inner core resembling the lamina propria, containing leukocytes, and mesenchymal and neuronal cells (Nash et al., 2021; Nash et al., 2023). In addition, a chicken 2D enteroid model that self-organises into an epithelial and mesenchymal sub-layer but lacks the underlying lamina propria cells has been developed (Orr et al., 2021). The aim of this study was to disentangle the innate immune response of epithelial cells and the immune response of the lamina propria cells to STm infection by comparing the gene expression profiles between uninfected and infected 2D and 3D enteroids. In addition, we analysed the effects of an invasion deficient strain, a Δ*prgH* mutant of STm, on the innate immune response in each enteroid model. Our study reveals marked differences in the response of intestinal epithelial cells to STm infection when lamina propria cells are present and therefore these models provide valuable insights into deciphering the distinct responses of epithelial and immune cells, such that findings with simpler cell-based models should be interpreted with caution.

## 1. Materials and Methods

### 2.1. Animals

Experiments were performed using embryonic day 18 (ED18) Hy-Line Brown fertile embryos (*Gallus gallus*) obtained from the National Avian Research Facility, University of Edinburgh, UK. Embryos were humanely culled under the authority of a UK Home Office Project Licence (PE263A4FA) in accordance with the guidelines and regulations of the Animals (Scientific Procedures) Act 1986.

### 2.2. Generation of chicken 2D and 3D enteroids

Tissue from duodenum, jejunum and ileum of ED18 chickens were retrieved and placed in phosphate buffered saline (PBS, Mg^2+^ & Ca^2+^ free) until use. For each independent culture, the intestines from five embryos were pooled. For the generation of 3D enteroids, the villi were released from the tissue as previously described (Nash et al., 2021). In brief, intestinal tissue was cut open longitudinally and cut into 3 mm pieces. The tissues were digested with *Clostridium histolyticum* type IA collagenase (0.2 mg/mL, Merck, Gillingham, UK) at 37°C for 50 min with shaking at 200 rpm. Single cells were removed by filtering the digestion solution through a 70 µM cell strainer (Corning, Loughborough, UK). The villi were obtained by rinsing the inverted strainer. The collected villi were centrifuged at 100 *g* for 4 min. The 3D enteroids were seeded at 200 villi per well in 24 well plates with 400 µl of Floating Organoid Media (FOM media); Advanced DMEM/F12 supplemented with 1X B27 Plus, 10 mM HEPES, 2 mM L-Glutamine and 50 U/mL Penicillin/Streptomycin (ThermoFisher Scientific (TFS), Paisley, UK).

For 2D enteroid generation, freshly isolated intestinal villi were enzymatically digested with Accutase (TFS) as previously described (Orr et al., 2021). To remove the majority of the fibroblasts, cells were resuspended in FOM media supplemented with 1X N2 supplement (TFS), 100 ng/mL human (hu) epidermal growth factor (huEGF, TFS), 10 µM CHIR 99021 (Stratech Scientific), 10 µM Y27632 (Stem Cell Technologies) and 100 nM LDN193189 (Cambridge Bioscience). Cells were incubated for 3 h at 37°C, 5% CO_2_ in 6 well plates. Non-adherent cells were removed, counted and seeded at 2X10^5^ cells in uncoated 24 well plates with 350 µl of FOM media supplemented with 1X N2 supplement, 100 ng/mL huEGF, 100 ng/mL huR-spondin, 50 ng/mL huNoggin (R&D Systems), 10 µM CHIR 99021 and incubated at 37°C, 5% CO_2_.

### 2.3. Preparation of *Salmonella*

*S.*Typhimurium strain ST4/74 nal^R^ (WT) is known to colonise the chicken intestine proficiently (Chaudhuri et al., 2013) and was routinely cultured in Luria-Bertani broth containing 20 µg/mL of naladixic acid (TFS). An isogenic ST4/74 nal^R^ Δ*prgH::kan* mutant, deficient in bacterial invasion, was additionally cultured in the presence of 20 µg/mL of kanamycin (Merck) and has been described previously (Balic et al., 2019). Both strains were transformed with a plasmid that constitutively expresses green fluorescent protein (GFP), pFVP25.1 (Valdivia and Falkow, 1996; Vohra et al., 2019), which was maintained in the presence of 50 µg/mL of ampicillin (Merck). Bacteria were incubated for 18 h at 37°C with shaking at 180 rpm to an optical density of 1 at 600 nm and pelleted at 2000 *g* for 10 min. Bacteria were washed twice with PBS and resuspended in 10 mL of PBS. Ten-fold serial dilutions were plated in duplicate on LB agar containing 20 µg/mL of naladixic acid incubated at 37°C overnight to determine viable counts retrospectively.

### 2.4. Bacterial infection and LPS treatment of 2D and 3D enteroids

On day 2 of culture, 3D enteroids were pelleted at 100 *g* for 4 min and reseeded at 200 enteroids per well on 24 well plates (Corning) in 400 µL of FOM media without antibiotics. Similarly, on day 2 of culture, 2D enteroids were washed twice with PBS and cultured for a further 24 h in FOM without antibiotics, CHIR and Y27632. After 24 h, on day 3 of culture, the 3D and 2D enteroids were treated with WT or Δ*prgH* STm strains (2X10^5^), LPS derived from STm (product code L6143, 1 µg/mL, Merck) or media only. At 4 and 8 hours post-infection (hpi) the supernatant was removed and cells washed with PBS and lysed in RLT Plus buffer (Qiagen) containing 10 µg/mL 2-mercaptoethanol (Merck). For increasing bacterial dose analysis, 3D enteroids were infected with 1X10^3^, 500 or 250 CFU of WT STm for 3 h. The 3D enteroids were further homogenised using QIAshredder columns (Qiagen). Samples were stored at -20°C until use.

### 2.5. Immuno-fluorescent staining and microscopy

For immuno-fluorescent staining, chicken 2D enteroids were grown on 2% Matrigel (Corning) coated transwell inserts (VWR, 0.33 cm^2^) in 24 well plates (Orr et al., 2021) while 3D enteroids were grown as outlined above. Chicken 2D and 3D enteroids were treated with WT or Δ*prgH* STm, LPS or media alone as outlined above. At 4 and 8 h post-treatment, cells were gently washed with PBS and fixed with 4% w/v paraformaldehyde (TFS) for 15 min at room temperature and blocked with 5% v/v goat serum in permeabilisation buffer (0.5% w/v bovine serum albumin and 0.1% w/v Saponin in PBS; Sigma-Aldrich). Permeabilisation buffer was used to dilute all antibodies. Cells were stained with mouse anti-human ZO-1 (Abcam, IgG1, clone A12) overnight at 4°C followed by the secondary antibody, goat anti-mouse IgG1 Alexa Fluor^®^594 (TFS) for 2 h on ice. Cells were counterstained with Hoechst 33258 and Phalloidin Alexa Fluor^®^647 (TFS) to stain for nuclei and F-actin, respectively. Slides were mounted using ProLong^TM^ Diamond Antifade medium (TFS). Controls comprising of secondary antibody alone were prepared for each sample. Images and Z-stacks were captured using an inverted LSM880 (Zeiss) with 40X and 63X oil lenses using ZEN 2012 (Black Edition) software and were analysed using ZEN 2012 (Blue Edition). Z-stack modelling was performed using IMARIS software (V9).

### 2.6. Isolation of RNA and reverse transcription

Total RNA from the enteroids was extracted using an RNeasy Plus Mini Kit (Qiagen) consisting of a genomic DNA column eliminator according to manufacturer’s instructions and quantified spectrophotometrically. Five independent 3D enteroids samples and three independent 2D enteroids samples that were of high quality were used for qPCR analysis (RNA concentration of >100 ng and a 260/230 ratio of 2). Reverse transcription was performed using the High Capacity Reverse Transcription Kit (Applied Biosystems) according to manufacturer’s instructions with random hexamers and oligo (dT)18, containing 100 ng of total RNA. The cDNA samples were stored in -20°C until use.

### 2.7. Pre-amplification and quantitative PCR using 96.96 Integrated Fluid Circuits dynamic array

Pre-amplification of cDNA was performed as previously described (Borowska et al., 2019; Bryson et al., 2023). In brief, 2.5 µl of a 200 nM stock pool of each primer pair (Supplementary File 1) was added to 5 µl of TaqMan PreAmp Master Mix (Applied Biosystems) and 2.5 µl of 1:5 dilution of cDNA. Due to its high level of expression, the ribosomal 28S (r28S) primer pair were excluded from the stock primer mix. Samples were incubated at 95°C for 10 min followed by 14 cycles of 95°C for 15 sec and 60°C for 4 min. Unincorporated primers were digested from the pre-amplified samples using 16 U/μl Exonuclease I (*E. coli*, New England Biolabs) at 37°C for 30 min. High-throughput qPCR was performed with the microfluidic 96.96 Dynamic array (Standard BioTools UK) as previously described (Borowska et al., 2019; Bryson et al., 2023). Each sample was run in duplicate with 88 target genes and 5 reference gene primers. In order to reduce inter-plate variation, an inter-plate calibrator (IPC) sample was employed on each array. The IPC sample comprised of pre-amplified cDNA derived from splenocytes stimulated with Concanavalin A (10 µg/mL, Sigma-Aldrich) for 4 h. Quantitative PCR was performed on the BioMark HD instrument (Fluidigm) using the thermal cycling conditions as previously reported (Borowska et al., 2019). The fluorescence emission was recorded after each cycling step. Raw qPCR data quality threshold was set to 0.65-baseline correction to linear (derivative) and quantitation cycle (Cq) threshold method to auto (global) using the Real-Time PCR Analysis software 3.1.3 (Fluidigm).

### 2.8. RT-qPCR data analysis

Data pre-processing, normalisation, relative quantification and statistics were performed using GenEx6 and GenEx Enterprise (MultiD Analyses AB). The qPCR performance baseline correction and set threshold of the instrument was compensated across the two array runs using the IPC samples. Delta Ct values were obtained by normalising the Ct values of the target genes with the geometric mean of three reference genes, GAPDH, TBP and r28S, identified from a panel of five references. The technical repeats were averaged and relative quantities were set to the maximum Cq value for a given gene. Relative quantities were log transformed (log2) and differentially expressed genes (DEG) between uninfected and infected enteroids was calculated by the −2^−ΔΔCT^ method (Livak and Schmittgen, 2001). Principal component analysis was performed using *prcomp* function and plots were generated using *ggplot* in RStudio (V1.1.442).

### 2.9. Statistical analysis

Statistical analysis of the differentially expressed genes (DEGs) between control (untreated) and LPS treated or between control and STm infected 2D or 3D enteroids was performed using GENEx5 and GenEx Enterprise (MultiD Analyses AB). Correction for multiple testing was performed with the estimation of false discovery rate using Dunn-Bonferroni correction threshold of 0.00054. After correction, DEG with a significant difference (*P*<0.05) and a fold change of ≥1.5 were identified. Finally, for each comparison, the fold change of untreated and treated samples at their respective timepoint was calculated. Graphs were prepared using GraphPad Prism 9.

## 3. Results

### 3.1. *Salmonella* Typhimurium alters tight junctions in chicken enteroids

We studied the early response of chicken 2D and 3D enteroid cultures to infection with STm in order to understand the relative contribution of epithelial and lamina propria cells to the response. In untreated and LPS-treated 2D and 3D enteroids actin expression was evenly distributed between cells and at the apical surface (Figures 1A & B). In chicken 2D enteroids infected with WT STm, raised F-actin structures were observed around the bacteria at the apical surface at 4 hpi (Figure 2A). Z-stack modelling of 2D enteroids demonstrates punctuated F-actin expression on the apical surface of a WT STm infected cell (Supplementary Video 1). At 8 hpi, bacteria could be detected within individual cells in 2D enteroids (Figure 2A). Despite the same inoculum used in WT STm infected 3D enteroids, we observed more remodelled F-actin around invading bacteria (Figure 2A). In contrast, no actin remodelling was observed in Δ*prgH* STm infected 2D and 3D enteroids although on occasion, bacteria were observed in close proximity to the apical surface of epithelial cells (Figure 2B) or found trapped in the folds of the epithelial buds (Supplementary Figure 1). This is consistent with the known role of T3SS-1 in promoting membrane ruffling and invasion.

**Figure 1.**
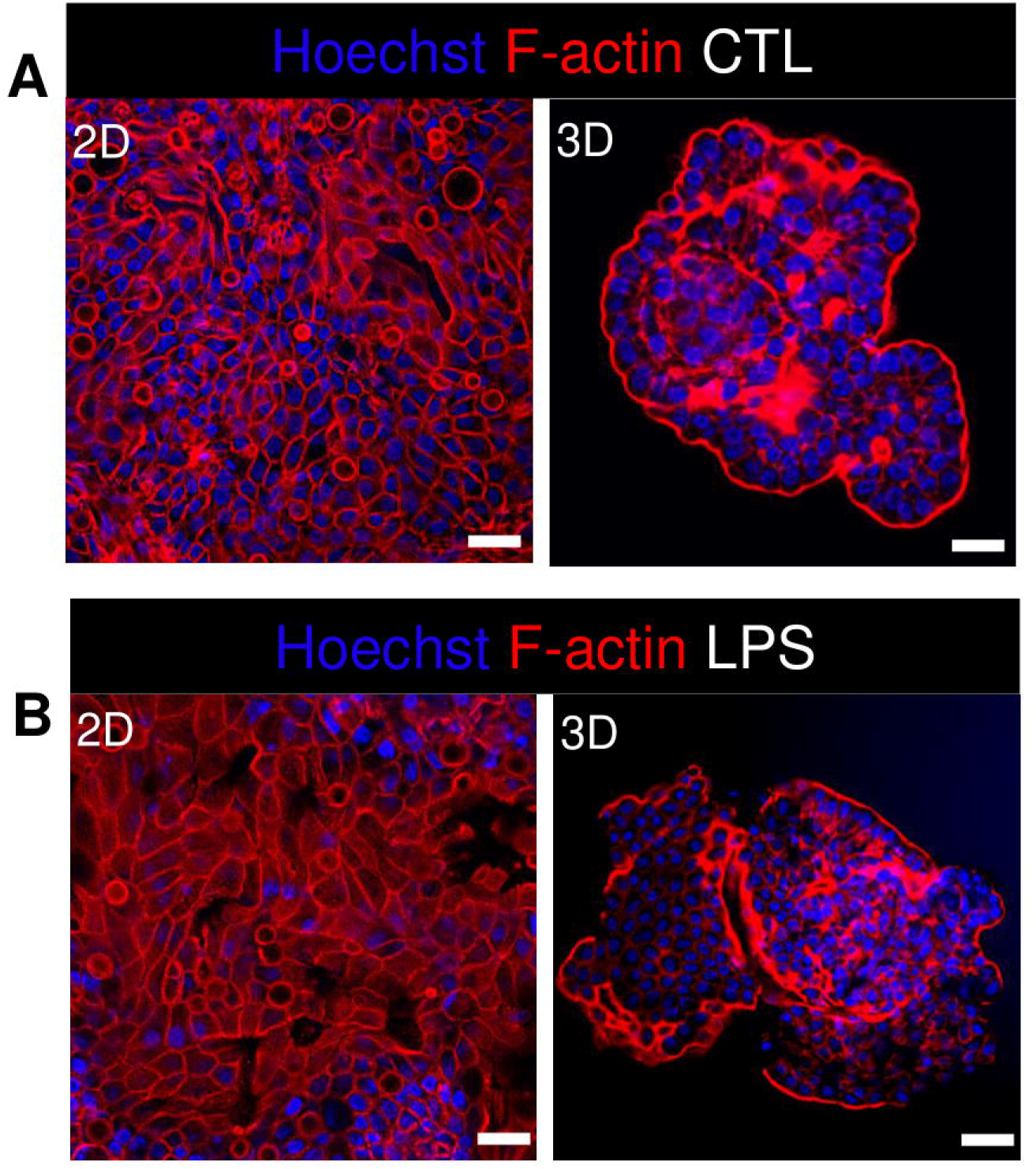
Untreated and LPS-treated 2D and 3D enteroids exhibit unaltered organisation of F-actin. Confocal micrographs of F-actin organisation in **A)** untreated or **B)** LPS-treated 2D and 3D enteroids. Images are representative of 3 independent experiments. Scale bars = 100 µm.

**Figure 2.**
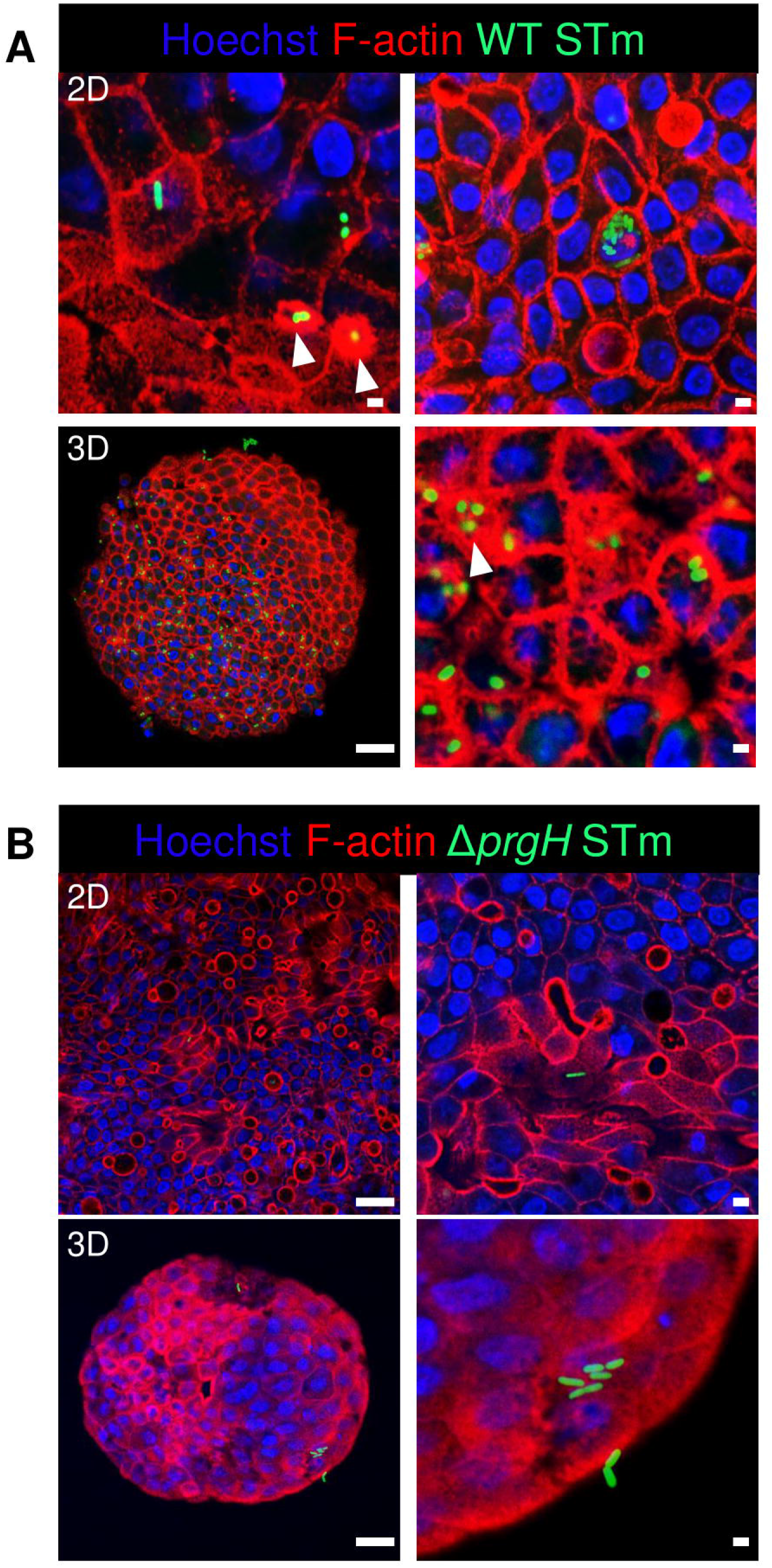
WT STm infection remodels F-actin in chicken enteroids. Confocal micrographs showing F-actin remodelling in **A)** WT STm infected 2D and 3D enteroids. At 8 hpi, dense F-actin staining can be observed surrounding the invading bacteria consistent with reorganisation of subcortical actin stimulating membrane ruffling (white arrows). The number of internalised bacteria was markedly higher in 3D enteroids inoculated with the same bacterial dose. **B)** No F-actin remodelling was observed in Δ*prgH* STm infected 2D and 3D enteroids, although bacteria were observed in close association with the apical surface of epithelial cells. Images are representative of 3 independent experiments. Scale bars = 100 µm and 50 µm.

Integrity of the epithelial barrier was analysed by immuno-fluorescent staining of the tight junction protein ZO-1. In untreated and LPS treated 2D and 3D enteroids, ZO-1 expression was localised to the lateral membrane, showing a typical polygonal shape of enterocytes and demonstrates that LPS had no effect on ZO-1 localisation (Figures 3A & B). In contrast, after treatment of 2D and 3D enteroids with WT STm, the localization of ZO-1 was discontinuous and jagged. In addition to being discontinuous, ZO-1 in 3D enteroids formed dense strands, which was not observed in 2D enteroids following infection. The Δ*prgH* STm strain did not elicit changes in ZO-1 distribution, which resembled that observed in the untreated 2D and 3D enteroids (Figure 4B). Occasionally, the Δ*prgH* STm strain was observed in close proximity to the apical surface of epithelial cells, but it did not cause any disruption to ZO-1 localisation (Supplementary Video 2).

**Figure 3.**
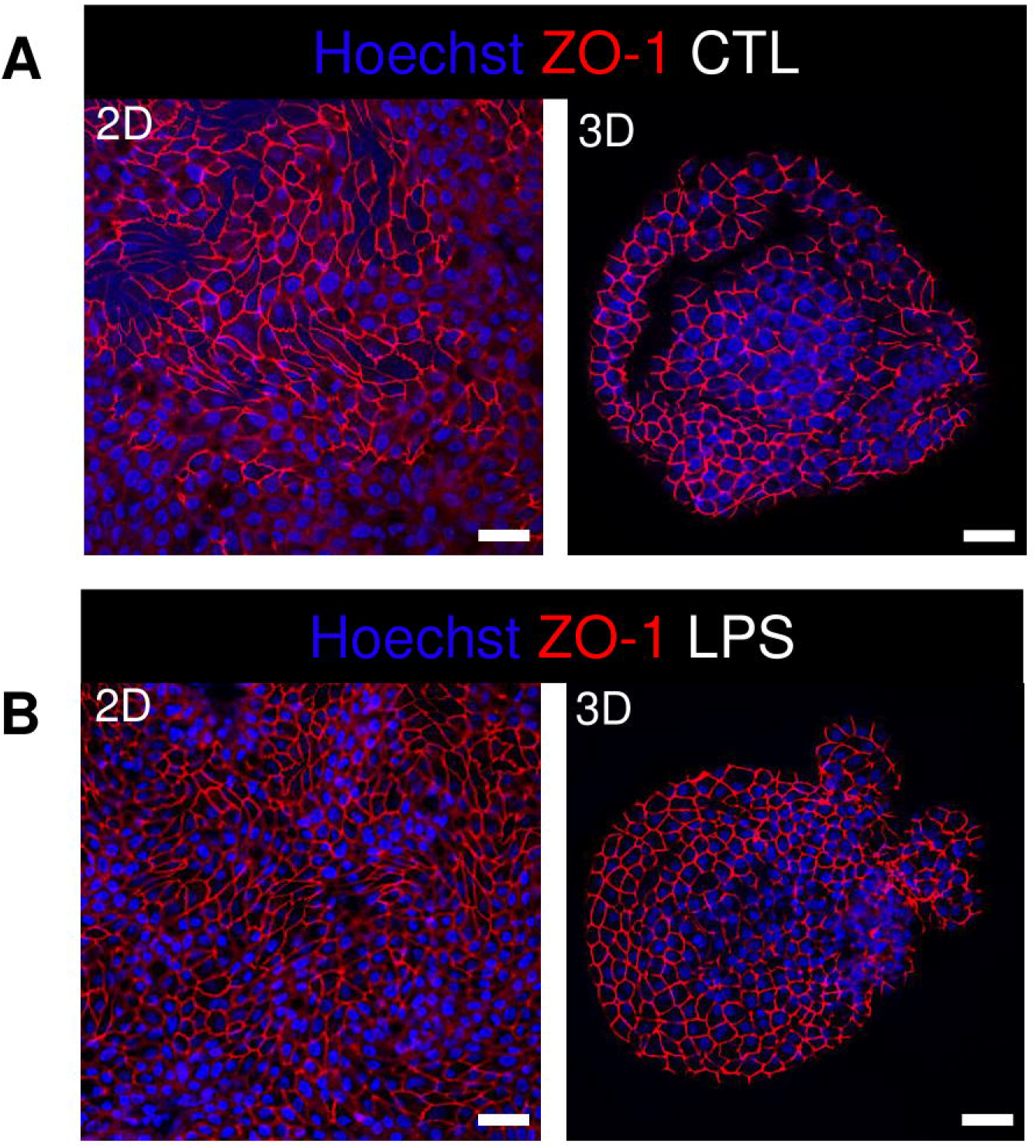
Distribution of the tight junction protein ZO-1 at cell-cell junctions in untreated and LPS treated enteroids. Confocal micrographs of ZO-1 distribution in **A)** untreated or **B)** LPS treated 2D and 3D enteroids show typical ZO-1 distribution at 8 h. Images are representative of 3 independent experiments. Scale bars = 100 µm.

**Figure 4.**
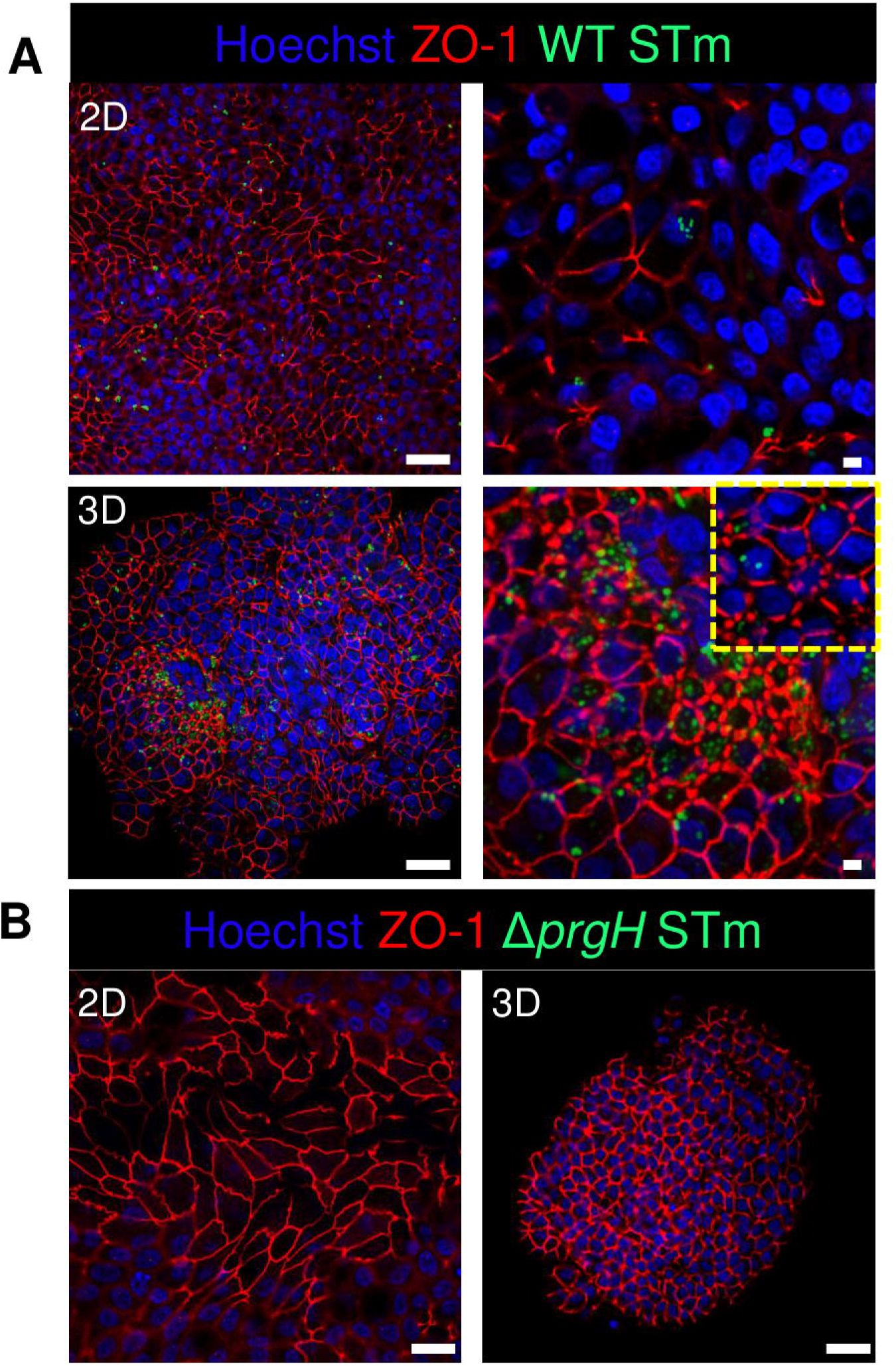
WT STm but not *ΔprgH* STm infection alters the distribution of the tight junction protein ZO-1 in chicken enteroids. Confocal micrographs showing the distribution of ZO-1 in **A)** WT STm infected 2D and 3D enteroids, which leads to reduced ZO-1 expression in 2D enteroids and altered ZO-1 distribution in 3D enteroids at 8 hpi (yellow dash insert image). **B)** ZO-1 distribution were unaltered in Δ*prgH* STm infected 2D and 3D enteroids at 8 hpi. Images are representative of 3 independent experiments. Scale bars = 100 µm and 50 µm.

### 3.2. Global transcriptional profiles cluster by treatment and culture model

To disentangle the innate immune responses of 2D and 3D chicken enteroids to *Salmonella* or its lipopolysaccharide, the mRNA expression levels of 88 innate-immune related genes were analysed using Fluidigm Biomark high-throughput qPCR at 4 and 8 h post-treatment. To assess the degree of heterogeneity between replicates and treatments, the global transcriptional profiles were compared using Principal Component Analysis (PCA). This analysis demonstrated that sample clustering was primarily by treatment as the uninfected and infected enteroids segregated from each other along the first principal component (PC1, Figures 5A & B). This difference accounted for 48-49% of the total variance in 2D and 3D enteroids respectively, and suggested that treatment is a greater determinant of transcriptional variance rather than time post-infection (PC2). PCA of the total dataset shows sample clustering based on the culture system, 2D versus 3D (Figure 5C, PC2), corresponding to the presence of different immune cell lineages in 3D enteroids while the effect of treatment (PC1) was preserved.

**Figure 5.**
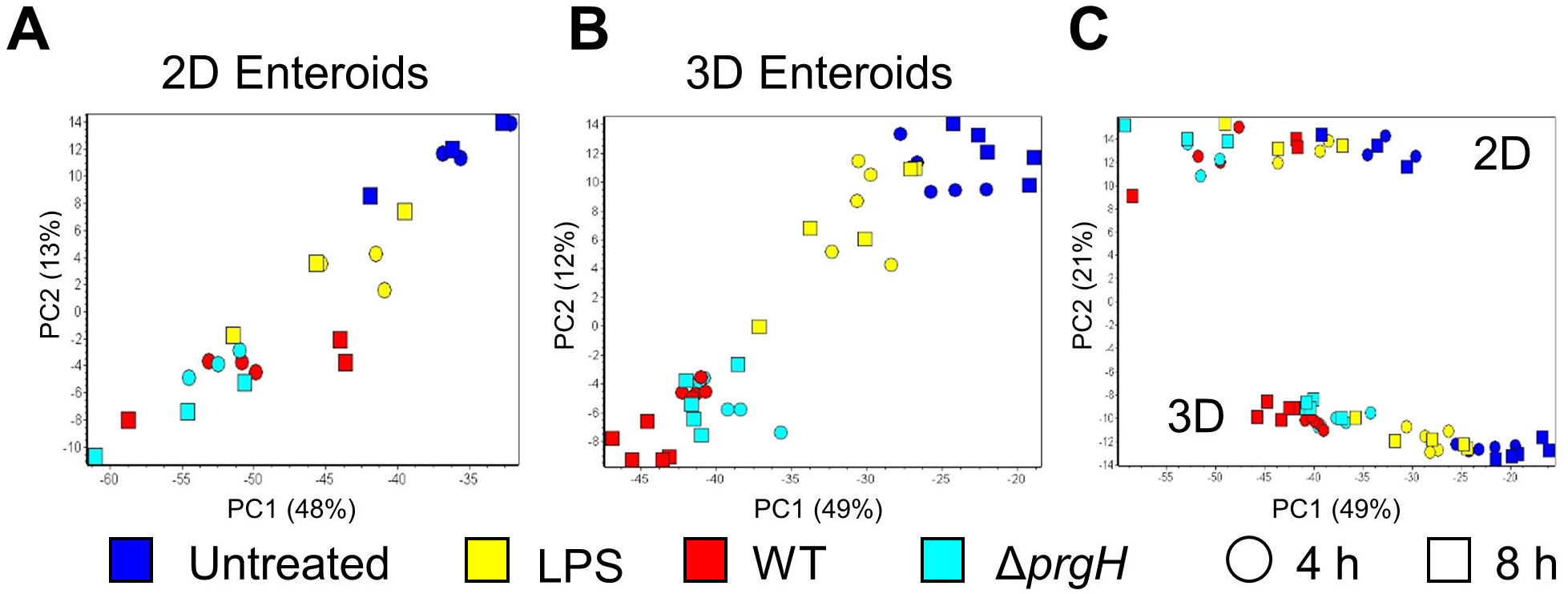
Principal component analysis (PCA) of gene expression profiles. PCA analysis of untreated, LPS treated, and WT and Δ*prgH* STm infected **A)** 2D enteroids (*n*=3) and **B)** 3D enteroids (*n*=5) at 4 hpi and 8 hpi alone and **C)** with both datasets.

### 3.3. Chicken 2D enteroids exhibit a more pronounced innate immune response to *Salmonella* LPS than 3D enteroids

Chicken 2D and 3D enteroids were treated with STm-derived LPS for 4 and 8 h. Firstly, the number of statistically differentially expressed genes (DEGs) with a fold change ≥1.5 at *P*<0.05 compared to their respective time-matched controls was analysed (Figure 6A). The number of DEGs was higher at 4 and 8 h in the 2D compared to 3D enteroids and the number of DEG substantially decreased with time in the 3D enteroids compared to the 2D enteroids. Next, the commonality and difference in the genes regulated in 2D and 3D enteroid were compared (Figure 6B). There was no core set of common DEGs between 2D and 3D enteroids at 4 and 8 h post-LPS treatment. Of the 11 DEGs regulated by 3D enteroids at 4 h post-LPS treatment, four were common to 2D enteroids, *C3ORF52*, *TNFAIP3*, *EAF2*, and *SDC4*. DEGs specifically regulated in 2D enteroids were related to TLR signalling (*TOLLIP, TRAF3IP2, TLR15*), effector protein functions (*LYZ*, *LYG2*) and regulation of immune responses (*BATF3, CD200L, IRF9, IL17REL, SOCS3*).

**Figure 6.**
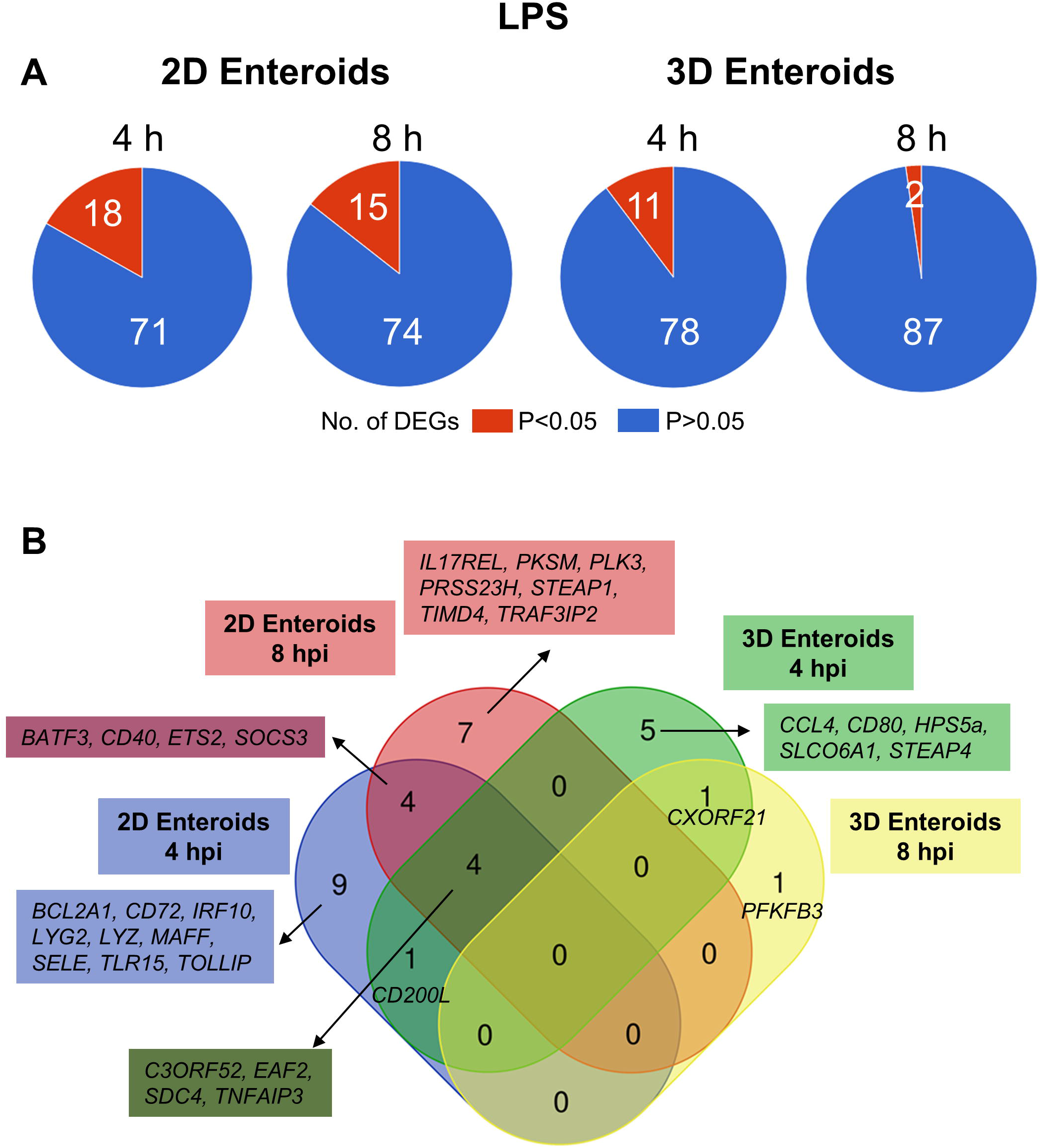
Chicken 2D enteroids exhibit more pronounced responses to LPS treatment. **A)** Pie charts indicating the number of statistically significant (*P*<0.05) and non-significant (*P*>0.05) DEGs in LPS treated 2D enteroids (*n*=3) and 3D enteroids (*n*=5) at 4 h and 8 h compared to time-matched, respective controls. DEGs with a significant difference (*P*<0.05) and a fold change of ≥1.5 were identified. **B)** Venn diagram showing common and unique DEGs across each enteroid model and timepoint.

The mRNA expression levels of a majority of the DEGs increased by 2-7 fold at *P* values of greater than 0.05 in LPS treated enteroids compared to untreated enteroids, except for *SELE* and *TIMD4,* which decreased in expression in 2D enteroids (Supplementary File 2). In conclusion, *Salmonella*-derived LPS treatment of chicken 2D and 3D enteroids resulted in differential innate immune responses, with 2D enteroids having a more pronounced response compared to 3D enteroids. These differences may suggest a regulatory role of lamina propria leukocytes to LPS exposure or suggest that the apical-out conformation of 3D enteroids inhibits TLR ligation at the basolateral region of epithelial cells whereas the TLRs expressed in a 2D layer may be more accessible.

### 3.4. Lamina propria leukocytes temporally govern the response to *Salmonella* infection

Next, the statistically significant DEGs (fold change ≥1.5, *P*<0.05 compared to their respective time-matched controls) was analysed in 2D and 3D enteroids infected with WT STm at 4 and 8 hpi. STm infected 2D enteroids differentially expressed 51 genes at 4 hpi, decreasing to 34 genes by 8 hpi (Figure 7A). In contrast, the number of DEGs in WT STm infected 3D enteroids was 54 at 4 hpi rising to 70 at 8 hpi (Figure 7A). There were 23 DEGs in common between 2D and 3D enteroids at both time-points (Figure 7B). This core set of genes are involved in the regulation of immune responses, *ATF3*, *BATF3*, *ETS2*, *IRF10*, *PTGS2, MAFA, NFκB2, TNFAIP3*, effector functions, *CCL4*, *CD200L*, *CD40*, *CD72*, *CD80*, *IL18* and TLR signalling, *EAF2, TLR15, TOLLIP.* The second largest set of common genes (16 DEGs) was shared between 2D enteroids at 4 hpi and 3D enteroids at 4 and 8 hpi. These genes are involved in regulating immune responses (*JUN, MAFF, PPARG, RASD1, SOCS1),* TLR/IL1β signalling *(MyD88, IL1R2),* and effector functions (*LYG2*, *TLR4)* while *TLR4* and the chemokine, *CCL5,* were differentially expressed at 4 hpi in both models.

**Figure 7.**
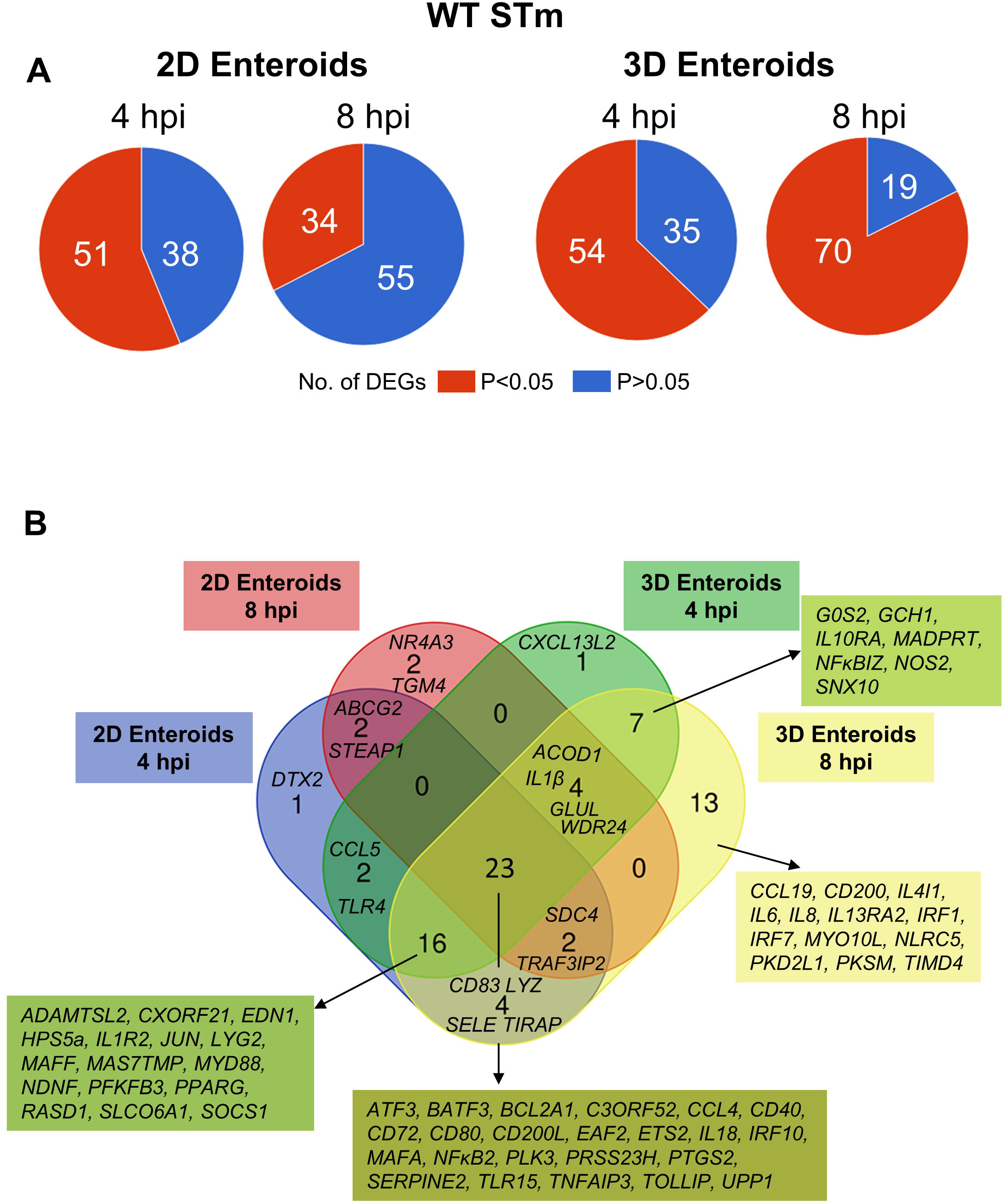
Innate immune response increase with time in WT STm infected 3D enteroids. **A)** Pie charts indicating number of statistically significant, (*P*<0.05) and non-significant (*P*>0.05) DEGs in WT STm infected 2D enteroids (*n*=3) and 3D enteroids (*n*=5) at 4 hpi and 8 hpi compared to their time-matched, respective controls. DEGs with a significant difference (*P*<0.05) and a fold change of ≥1.5 were identified. **B)** Venn diagram showing common and unique DEGs across each enteroid model and timepoint.

The number of DEGs that were uniquely expressed in one or the other model differed substantially, with the 2D enteroids expressing 5 unique genes; specifically those encoding a regulator of Notch signalling (*DTX2)*, a membrane transporter (*ABCG2*), a metalloreductase (*STEAP1*), a nuclear receptor (*NR4A3*), and transglutaminase (*TGM4*). In contrast, the 3D enteroids uniquely expressed 21 genes upon STm infection with a majority associated with immune cell functions. The seven genes shared between 4 and 8 hpi are involved in regulating inflammation (*IL10RA, IL13RA2, NFκBIZ*), bactericidal activity (*GCH1, NOS2*), membrane trafficking in endosomes (*SNX10*), and apoptosis (*GOS2*). By 8 hpi, genes involved in the regulation of interferon (IFN) and IFN-inducible genes (*IRF1*, *IRF7*, *NLRC5*) and macrophage and DC activities (*CCL19*, *CD200*, *IL4IL*, *TIMD4*) were found to be differentially expressed.

The temporal changes in the fold change levels of the 23 common DEGs demonstrates that the vast majority were expressed at higher levels in the 2D enteroids compared to the 3D enteroids at 4 hpi (Supplementary File 2, Figure 8A). In contrast, at 8 hpi the majority of the common genes had higher fold change levels in the 3D enteroids compared to 2D enteroids. The magnitude of the fold change for DEGs in 2D enteroids did not exceed 96 (*C3ORF52*), whereas in 3D enteroids three common DEGs, possibly involved in inhibiting inflammatory responses, were expressed at a fold change ranging from 114-905 (*CD72*, *CD80, TNFAIP3*). In addition, the magnitude of the fold change of all 23 common genes increased with time in 3D enteroids whereas 13 of the common genes decrease with time in 2D enteroids (Figure 7B). Overall, the data suggests that lamina propria leukocytes, present in 3D enteroids, temporally govern the expression levels of a core gene set in response to STm infection, through the unique expression of genes associated with immune cell function regulating inflammation.

**Figure 8.**
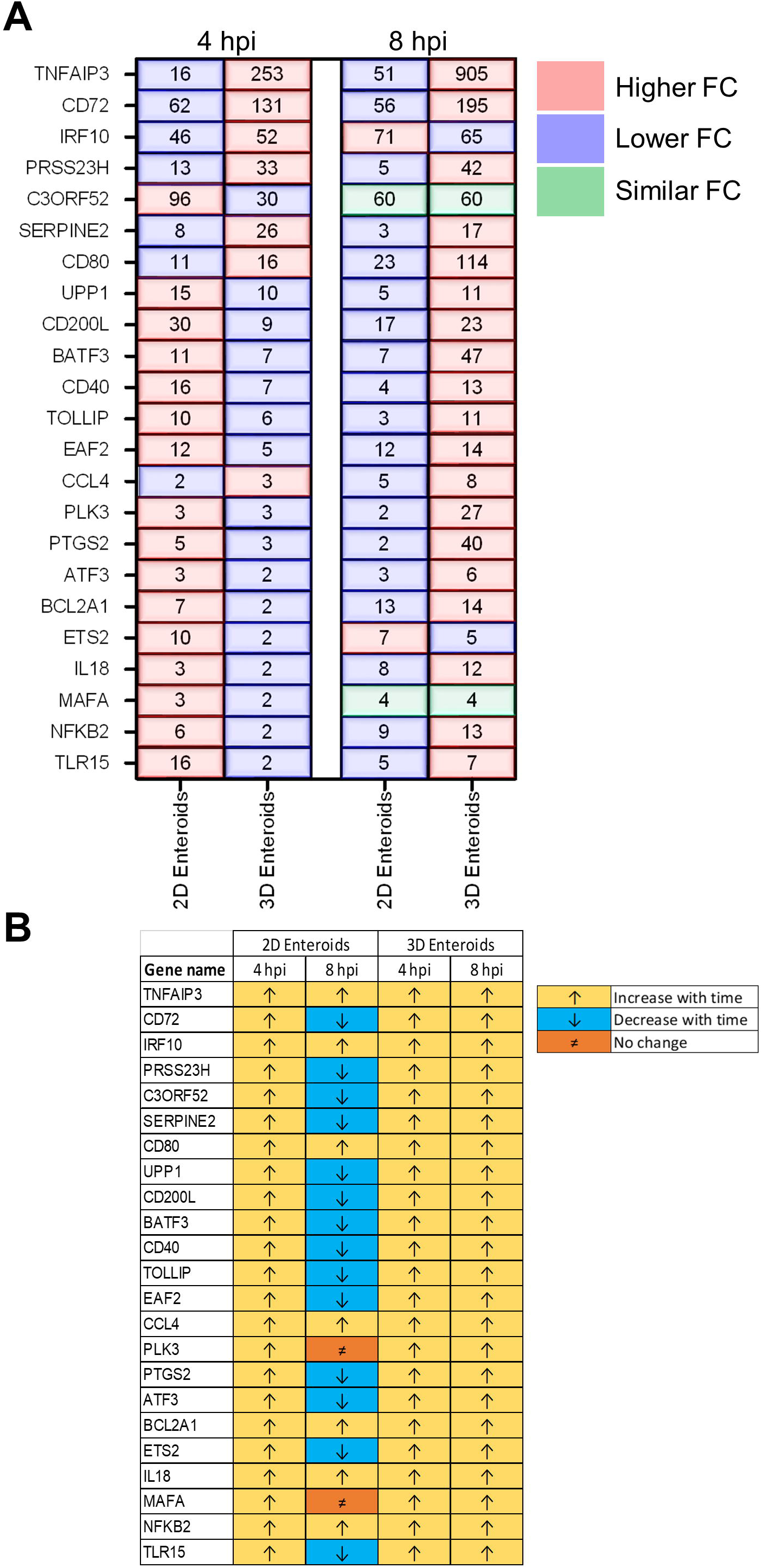
Lamina propria leukocytes temporally govern the response to WT STm infection. **A)** A comparison of the fold change levels of the common DEGs between the enteroid models demonstrates increased expression levels in 2D enteroids compared to 3D enteroids at 4 hpi. At 8 hpi, the expression levels of the common DEG are higher in 3D enteroids compared to 2D enteroids. **B)** The expression levels of a majority of the common DEGs decrease with time in WT STm infected 2D enteroids and increase with time in 3D enteroids. Fold change values represent the mean of 3 (2D) or 5 (3D) independent experiments, relative to their time-matched respective controls.

### 3.5. Chicken enteroids mount an innate immune response to non-invasive *Salmonella*

To characterise the effects of a non-invasive STm strain on innate immune responses, 2D and 3D enteroids were infected with the isogenic Δ*prgH* mutant of STm strain 4/74. At 4 and 8 hpi, 56 and 39 genes were differentially expressed in 2D enteroids at *P* values <0.05, fold change of ≥1.5, respectively (Figure 9A). In 3D enteroids, infections with the Δ*prgH* STm strain differentially affected 52 genes at 4 hpi at fold change of >2 and *P* values <0.05, increasing to 68 genes by 8 hpi (Figure 9A). Analysis of the DEGs across the models and time-points indicated a core set of 28 common genes (Figure 9B). Similar to WT STm infection, genes involved in the regulation of immune responses (*ATF3*, *BATF3*, *ETS2*, *IRF9*, *NFκB2, MAFA, TNFAIP3*), effector functions, (*CCL4*, *CD200L*, *CD40*, *CD72*, *CD80*, *IL18)* and TLR signalling, (*EAF2, TLR15, TRAF3IP2, TOLLIP*) were upregulated after *ΔprgH* STm infection in both models (Supplementary File 2). An additional four genes were upregulated in *ΔprgH* STm infection and were involved in the regulation of glycolysis (*PFKFB3*) and cytokine signalling (*SOCS3*), and intracellular signalling (*RASD1* and *TRAF3IP2*). When disentangling the innate responses between the models, 8 and 20 genes were differentially expressed only in 2D and 3D enteroids, respectively. Similar to the response to WT STm infection, Δ*prgH* STm infected 2D enteroids regulated *CCL5* and *DTX2* at 4 hpi and genes involved in the regulation of antigen presentation (*CD83*), and inhibition of NFκB activation (*NFAIP2*). At both time-points, a gene involved in cell-cell junctions (*STEAP1*) and a serine protease (*PRSS23*) were upregulated and by 8 hpi, only one DEG, *TIMD4*, encoding a phosphatidylserine receptor for apoptotic cells, was specifically regulated in 2D enteroids. The shared genes in 3D enteroids were involved in nitric oxide synthesis (*GCH1, NOS2*), negative regulation of inflammation (*NFκBIZ*, *SOCS1*), the mTOR signalling pathway (*WDR24*), and membrane trafficking in endosomes (*SNX10*).

**Figure 9.**
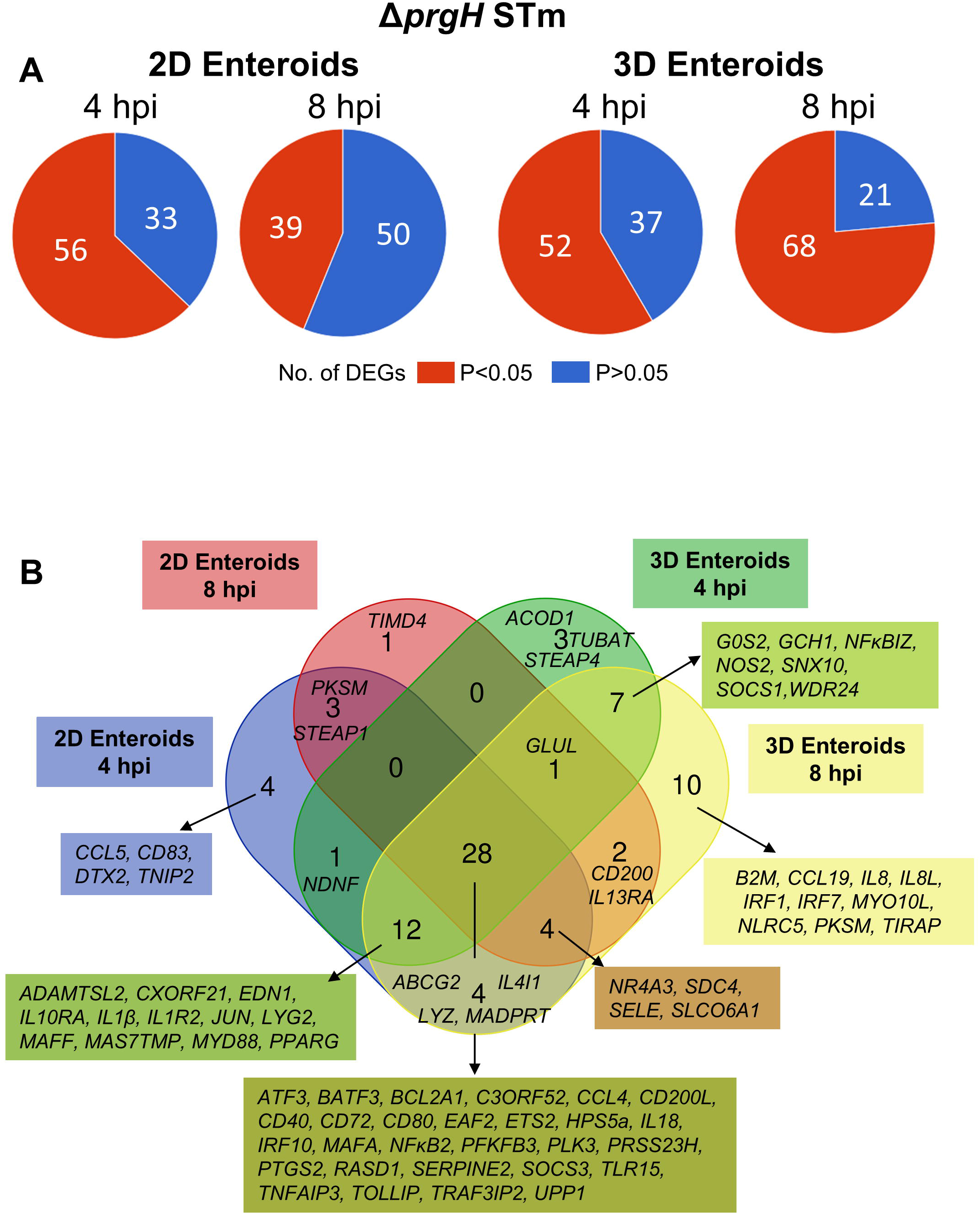
Chicken enteroids mount an innate immune response to non-invasive STm. **A)** Pie charts indicating the number of statistically significant (*P*<0.05) and non-significant (*P*>0.05) DEGs in STm infected 2D enteroids (*n=3*) and 3D enteroids (*n=5*) at 4 hpi and 8 hpi, compared to their time-matched, respective controls. DEGs with a significant difference (*P*<0.05) and a fold change of ≥1.5 were identified. **B)** Venn diagram showing common and unique DEGs across each enteroid model and timepoint.

Fold change comparisons of the common genes demonstrates the expression levels for most genes were higher in 2D enteroids compared to 3D enteroids at 4 hpi (Figure 10A). However, by 8 hpi the expression levels for 15 out 28 common DEGs were higher in 2D enteroids compared to 3D enteroids. In contrast to infection with WT STm, when a differential response was detected between 2D and 3D enteroids over time, treatment with the invasive deficient Δ*prgH* STm strain resulted in the downregulation of almost half of the common DEGs in both 2D and 3D enteroids (Figure 9B). The data indicates that both the chicken 2D and 3D enteroids exhibit distinct innate immune responses, which may be attributed to the presence of lamina propria cells.

**Figure 10.**
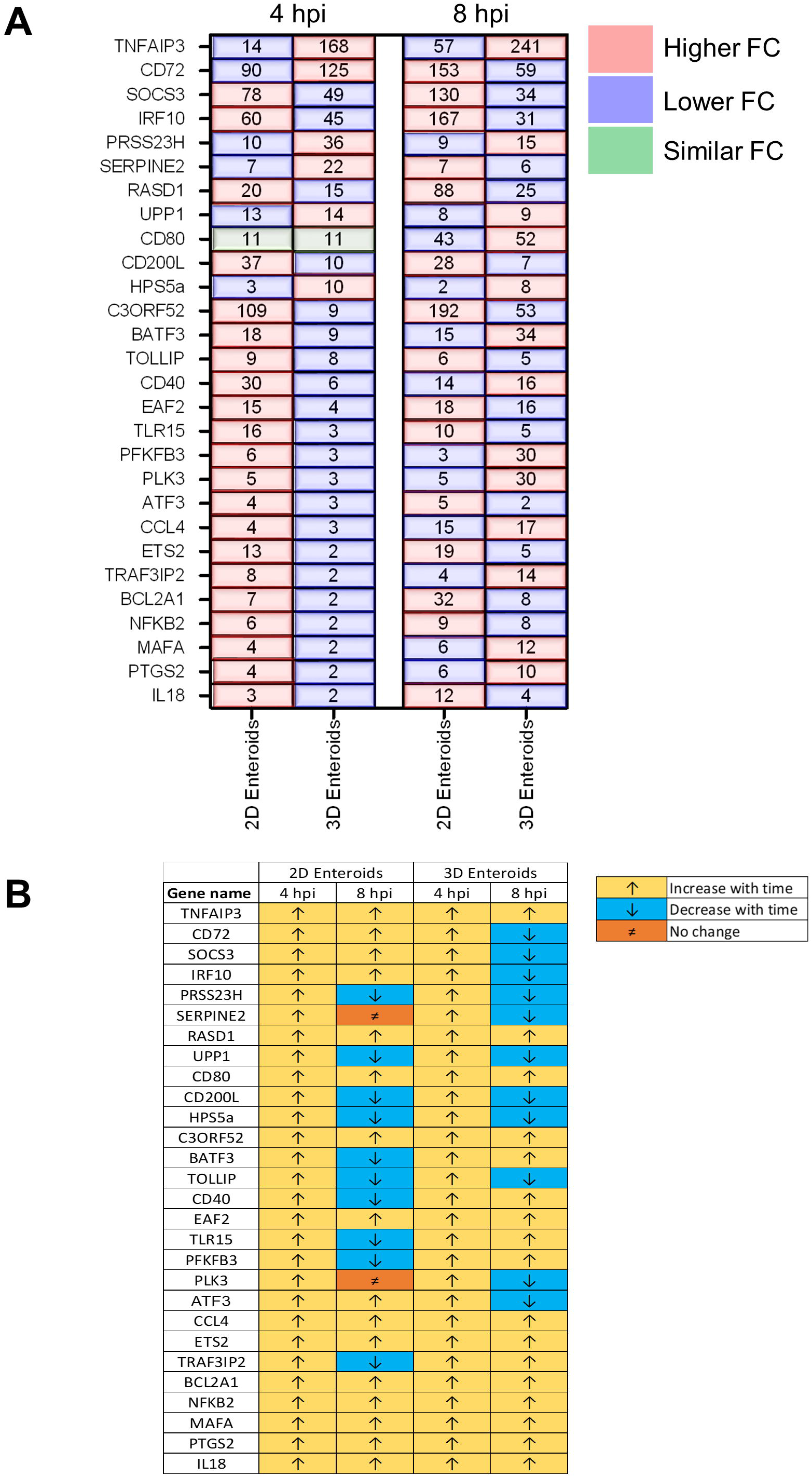
*ΔprgH* STm induces a rapid but brief innate immune response in chicken enteroids. **A)** Fold change difference in the common DEGs in Δ*prgH* STm infected 2D and 3D enteroids at 4 hpi and 8 hpi. **B)** The innate immune responses decrease with time in Δ*prgH* STm infected 2D and 3D enteroids. Fold change values represent the mean of 3 (2D) or 5 (3D) independent experiments, relative to their time-matched respective controls

### 3.6. Transcriptional regulation of innate immune genes remain unaffected by bacterial invasion

When comparing the innate immune responses between 2D and 3D enteroids, a majority of the DEGs were common to both models when infected with the invasive WT STm or non-invasive Δ*prgH* STm. We next analysed the contribution of the bacterial load to innate responses by comparing the DEGs (fold change of ≥1.5 and *P*<0.05) in WT and Δ*prgH* STm infected enteroids at the late stage of infection, 8 hpi, when we observed numerous cells infected with WT STm in both models. Irrespective of STm strain, 26 DEGs were common across each model demonstrating the transcriptional regulation of these genes is independent of bacterial invasion (Figure 11). Few DEGs were uniquely upregulated by either WT or Δ*prgH* STm, independent of the enteroid model. WT and Δ*prgH* STm infected 2D enteroids shared 1 common gene (*STEAP1*) and WT and Δ*prgH* STm each regulated 1 specific gene each, *TGM4* and *TLR4*. WT and Δ*prgH* STm infection of 3D enteroids shared 27 common genes while WT STm infection regulated 4 genes, (*CD83, IL6, NDNF, PKD2L1*) and *ΔprgH* STm regulated 3 genes (*ABCG2, B2M, IL8L*).

**Figure 11.**
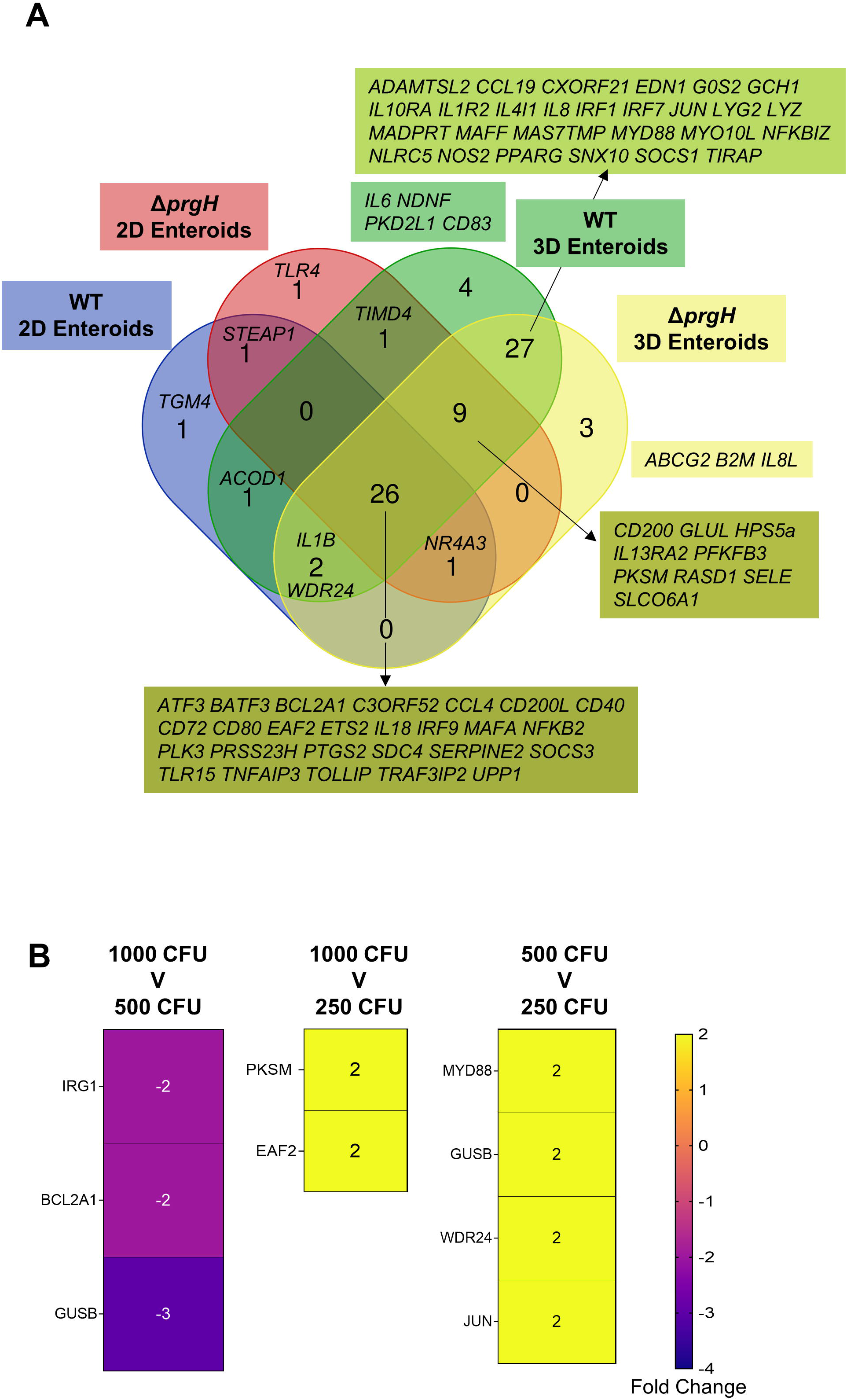
Transcriptional regulation of innate immune responses does not require bacterial invasion in chicken enteroids. **A)** Venn diagram showing the DEGs that are altered by WT or Δ*prgH* STm infected 2D and 3D enteroids at 4 hpi and 8 hpi. **B)** Heat maps of the fold change difference in the DEGs (Fold change ≥1.5, *P*<0.05) between 1000, 500 or 250 CFU of WT STm infected 3D enteroids demonstrates that the transcriptional regulation of innate immune genes is not significantly affected by bacterial burden. Fold change values represent the mean of 3 independent experiments.

To determine the effects of bacterial burden on the innate immune responses of lamina propria cells, we infected 3D enteroids with 1000, 500 and 250 CFU of WT STm for 3 h and analysed the mRNA expression levels using Fluidigm Biomark high-throughput qPCR. Few genes were significantly differentially regulated, with a fold change of ≥1.5 and *P* values <0.05, between the different doses of WT STm (Figure 11B). Moreover, the fold change of these DEGs was low. This demonstrates that the burden of bacteria does not significantly alter the transcriptional regulation of innate immune genes in chicken 3D enteroids and further supports the role lamina propria cells have on the regulation of responses to *Salmonella*.

## 4. Discussion

The aim of this study was to disentangle the contribution of avian intestinal epithelial cells and lamina propria cells to STm infection using unique enteroid culture models. In contrast to mammalian enteroids or intestinal organoids, the inner core of chicken 3D enteroids consists of cells present in a lamina propria including T and B cells, natural killer cells, macrophages, dendritic cells (DC) and heterophils (Nash et al., 2021; Nash et al., 2023), whereas few intraepithelial leukocytes are present in chicken 2D enteroids (Orr et al., 2021). Intestinal epithelial cells are equipped with pattern recognition receptors, such as TLRs, that initiate inflammatory responses in response to pathogen-associated molecular patterns if the epithelial barrier is breached. Additionally, intestinal epithelial cells play a crucial role in maintaining homeostasis and the unique mucosal immunological equilibrium [reviewed in (Mahapatro et al., 2021)]. The regulation of immune function and cross talk between intestinal epithelial cells and the underlying lamina propria cells is not well understood. The limited knowledge in this area in chickens can be partly attributed to the lack of cell lines or complex primary cell culture systems. Our 2D and 3D avian enteroids therefore offer a valuable model system in which to understand the contributions of lamina propria cells to the response to a model pathogen and its lipopolysaccharide as a major innate immune agonist.

Previously, we demonstrated that WT STm can adhere to chicken 3D enteroids and enter cells in a manner associated with actin remodelling (Nash et al., 2021). In this study, we used the same invasive WT STm and an isogenic invasion deficient Δ*prgH* mutant strain and demonstrated that the latter was adherent and immunogenic but did not invade chicken 2D and 3D enteroids. We observed that WT STm remodelled F-actin and disrupted tight junctions based on altered localisation of ZO-1, whereas these phenotypes were not seen with the Δ*prgH* STm strain or after treatment with *Salmonella* LPS. ZO-1 associates with tight junction proteins, claudins and occludins, required to maintain cellular polarity and the paracellular barrier (Odenwald et al., 2017). Our data are consistent with the known role of T3SS-1 and its effector proteins in *Salmonella* interactions with mammalian intestinal epithelial cells *in vitro* and *in vivo* (Jepson et al., 2000; Boyle et al., 2006; Lhocine et al., 2015).

The innate immune gene expression levels was analysed to investigate the common responses elicited by *Salmonella* in 2D and 3D enteroids. We found that each model modulates the expression of common genes in a temporal fashion. In WT STm infected 3D enteroids, expression levels of the common DEGs increased with time but in 2D enteroids, 60% of the common genes expression levels decreased with time. Twenty-eight DEGs were shared between Δ*prgH* STm infected 2D and 3D enteroids but in contrast to WT STm infection, the expression levels of 42% of the common DEGs decreased with time in 3D enteroids. While the Δ*prgH* STm strain was not observed to invade 2D or 3D enteroids, it could be found in close association with their apical surface. This association is likely mediated by bacterial adhesins such as fimbriae, flagella and outer membrane proteins [reviewed by (Stones and Krachler, 2016)], and may suffice to activate innate immune responses in enteroids. Comparison of the responses by invasive and non-invasive STm infected 3D enteroids show 91% of the DEGs are in common indicating that the upregulation of the genes measured in our study are independent of invasion and bacterial load. We can be confident that the differences in gene expression observed upon infection of 2D and 3D enteroids reflect the presence of lamina propria cells, as opposed to differences in the level of bacterial uptake, owing to the similarity of responses of enteroids to infection with WT or Δ*prgH* STm. However, the magnitude of the responses from the lamina propria cells are dependent on invasion as expression levels of all genes increased with time in WT infected and not Δ*prgH* STm infected 3D enteroids. The strong responses after treatment with Δ*prgH* STm may be related to the high susceptibility of day-old chicks to *Salmonella* compared to week-old chickens (Withanage et al., 2005). In addition, the ligand for flagella, TLR5, is highly expressed throughout the small intestine of neonatal mice (Price et al., 2018). Our study utilised enteroids derived from ED18 embryos, therefore it is possible that the effects observed with the STm mutant are related to the susceptibility of immunologically immature models and TLR5 expression during the neonatal period. Further research is necessary to understand the effect of age and the role of different virulence factors on the innate responses to STm.

A core set of pro-inflammatory related transcription factors were upregulated in the STm infected 2D and 3D models such as *ATF3*, *BATF3* and *MAFA*. These genes play vital roles in modulating glucose homeostasis, immunity, cell differentiation and cell function (Wu et al., 2021). The transcription factor IRF10, that is not present in humans and mice (Santhakumar et al., 2017), was also upregulated and is known to respond to viral and bacterial infections and regulate the expression of the IFN-γ target genes (Nehyba et al., 2002; Zhu et al., 2020). *Salmonella* has acquired mechanisms to survive and exploit the inflammatory response which in turn progresses infection and replication in the host (Gibbs et al., 2020). We found genes such as, *TOLLIP*, *RASD1* and *SOCS1,* which are involved in the negative regulation of downstream TLR4 signalling, NFĸB activation and cytokine activities, were upregulated across all models. Collectively, the modulation of a majority of the shared genes resemble the immune landscape similar to *in vivo* studies (Berndt et al., 2007; Fasina et al., 2008; Khan and Chousalkar, 2020; Bescucci et al., 2022; Dar et al., 2022). Overall, the data suggests a tight balance between pro- and anti-inflammatory responses is taking place in both 2D and 3D enteroids during STm infection.

Several genes exhibited enteroid model-specific differential expression in response to STm, related to the presence (3D) or absence (2D) of lamina propria cells. In 2D enteroids, an epithelial cell-specific response was evident. *DTX2* was upregulated in 2D enteroids which regulates the NOTCH signalling pathway, supporting epithelial cell homeostasis in terms of stem cell and epithelial cell maintenance (Artavanis-Tsakonas et al., 1999). Other DEGs that were uniquely upregulated in 2D enteroids include the metalloreductase, *STEAP1,* which is expressed at cell-cell junctions and facilities epithelial-mesenchymal transition and is involved in metal metabolism. *STEAP1* was also found to be upregulated in the liver of *S*. Enteritidis infected broiler chickens (Coble et al., 2013). *ABCG2* was also upregulated and is a membrane transporter gene of the superfamily of ATP-binding cassette (ABC) transporters. ABCG2 plays a protective role in blocking absorption at the apical membrane of the intestine (Vlaming et al., 2009). *NR4A3* is a nuclear receptor that contributes to apoptosis (Fedorova et al., 2019) and antibacterial autophagy (Sorbara et al., 2018). *TGM4* was upregulated in STm infected 2D enteroids which in chickens may have a similar function as in rat intestinal epithelial cells, e.g. cross-linking of immunoglobulin opsonised antigens to CD89 or Fc receptors on antigen-presenting cells (Rychlik et al., 2014). Together the significantly upregulated genes specific to chicken 2D enteroids are involved in a range of metabolic and transport processes associated with epithelial cell functions.

In 3D enteroids, we observed a pronounced response related to activation of macrophages and DC. Within macrophages and DC, STm can survive within *Salmonella*-containing vacuoles, including by inhibiting phagolysosome fusion and the respiratory burst (Buchmeier and Heffron, 1991). Key phagocyte chemoattractant genes, *IL8* (CXCLi1) and *IL8L* (CXCLi2) and transcription factors, *IRF1* and *IRF7,* were only upregulated in 3D enteroids (Poh et al., 2008). In mice, expression of *IRF1* in DC is required for Th17 generation during STm infection (Park et al., 2019) whereas TLR4 signalling in macrophages uniquely engages IRF1, facilitating access of the interferon stimulatory genes loci for transcription (Park et al., 2019). *IRF7* is also upregulated in chicken heterophils stimulated with *S*. Enteritidis leading to the induction of IFNs (Kogut et al., 2012). In addition, we observed the upregulation of *IL4l1*, a putative anti-inflammatory gene expressed by myeloid cells (Marquet et al., 2010) and one of the most inducible genes found in the chicken cecum post-*S*. Enteritidis infection (Elsheimer-Matulova et al., 2020). One gene, *NOS2*, was consistently downregulated in 3D enteroids irrespective of the STm strain. *NOS2*, expressed by macrophages, is supressed by *S*. Enteritidis in HD11 chicken macrophage cell line (He et al., 2012). In murine macrophages, STm induces IFN-γ and NOS2 production which in turn represses CCL3 and CCL4 production (Chandrasekar et al., 2013). In our studies, *CCL4* was upregulated across both models while *CXCL13L2* and *CCL19* were specifically upregulated in 3D enteroids. CCL19 is a key chemokine involved in DC trafficking and its cognate receptor, CCR7, is upregulated on murine DC upon uptake of STm (Cheminay et al., 2002). In line with the significant downregulation of *NOS2*, this gene expression pattern may suggest a typical reprogramming from M1 to M2 macrophages during infection in 3D enteroids. Although the M1-M2 dichotomy has not been established in chicken macrophages, WT STm is known to induce a low M1 and high M2 response in porcine alveolar macrophages which was dependent on the T3SS-1 (Kyrova et al., 2012). ScRNA-Seq profiling of *S*. Enteritidis infected murine macrophages demonstrated that the bacterium subvert cells towards a M2-like macrophage phenotype to prevent killing (Saliba et al., 2016). Further studies are warranted to disentangle the *Salmonella*-dependent M1/M2 dichotomy in chickens. Our study clearly indicates that 3D enteroids have an innate response to STm that is lacking in 2D enteroids which is associated with macrophages and DC.

## 5. Conclusion

In conclusion, the chicken 2D and 3D enteroids allowed for the first description of the distinct innate immune responses exhibited by epithelial cells and lamina propria leukocytes. The enteroid models successfully replicated several observations also demonstrated after *in vivo* infection of chickens with *Salmonella*, including the alteration of tight junctions and the induction of inflammatory responses. The 3D enteroid model offers many advantages over other models to reduce animal use in the study of host-pathogen interactions.

## Data availability statement

The original contributions presented in the study are included in the article/Supplementary material; further inquiries can be directed to the corresponding authors.

## Ethics statement

All procedures were conducted under Home Office project license PE263A4FA according to the requirements of the Animal (Scientific Procedures) Act 1986, with the approval of The Roslin Institute’s Animal Welfare and Ethical Review Board. Embryos were humanely culled in accordance with Schedule 1 of the Animals (Scientific Procedures) Act 1986.

## Author contributions

KS, TN and LV conceptualised the study; LV secured funding to undertake the work and supervised the project; KS, TN, SS, DB, and JM performed the experiments and analysed the data with support of PV, MS and LV. KS wrote the manuscript supported by SS, PV, MS and LV. All authors read and approved the final manuscript.

## Funding

This work was supported by the Biotechnology and Biological Sciences Research Council Institute Strategic Program Grant funding (BB/X010937/1, BBS/E/D/10002071, BBS/E/D/10002073 and BBS/E/D/20002174) and an iCase doctoral studentship to TN (BB/MO14819).

## Acknowledgements

We wish to thank the animal caretakers of the National Avian Research Facility for the supply of eggs and for helpful advice from staff at the Roslin Institute’s Bio-imaging Facility. For the purpose of open access, the author has applied a Creative Commons Attribution (CC BY) licence to any Author Accepted Manuscript version arising from this submission.

## Conflict of interest

The authors declare that the research was conducted in the absence of any commercial or financial relationships that could be construed as a potential conflict of interest.

## Supplementary material

Supplementary Table 1: Gene accession numbers and primer information

Supplementary Table 2: Fold change and *P* values levels of DEGs in WT STm infected or Δ*prgH* STm and STm LPS treated 2D and 3D enteroids at 4 and 8 hpi

Supplementary Video 1: Z-stack modelling of WT STm infected 2D enteroids at 8 hpi

Supplementary Figure 1: Z-stack modelling of Δ*prgH* STm infected 2D enteroids at 8 hpi

Supplementary Video 2: Confocal analysis of Δ*prgH* STm infected 3D enteroids at 8 hpi

## Notes

### Competing Interest Statement

The authors have declared no competing interest.

