## Supplementary Video 1 for "Disentangling the Innate Immune Responses of Intestinal Epithelial Cells and Lamina Propria Cells to *Salmonella* Typhimurium Infection in Chickens"

### Slide 1
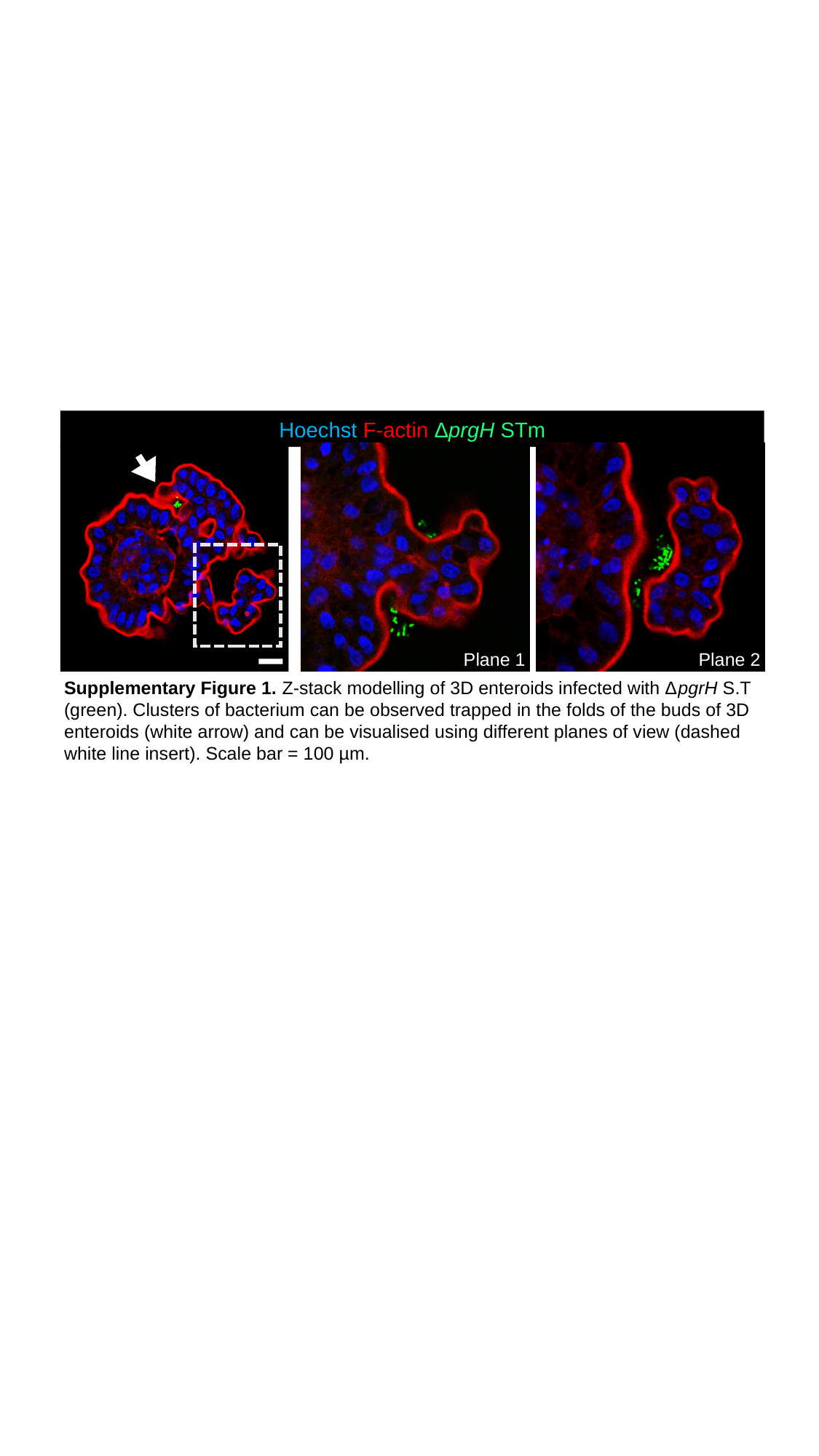

Hoechst F-actin ΔprgH STm
Plane 1
Plane 2
Supplementary Figure 1. Z-stack modelling of 3D enteroids infected with ΔpgrH S.T (green). Clusters of bacterium can be observed trapped in the folds of the buds of 3D enteroids (white arrow) and can be visualised using different planes of view (dashed white line insert). Scale bar = 100 µm.
