## Supplementary Figure 1 for "Disentangling the Innate Immune Responses of Intestinal Epithelial Cells and Lamina Propria Cells to *Salmonella* Typhimurium Infection in Chickens"

### Slide 1
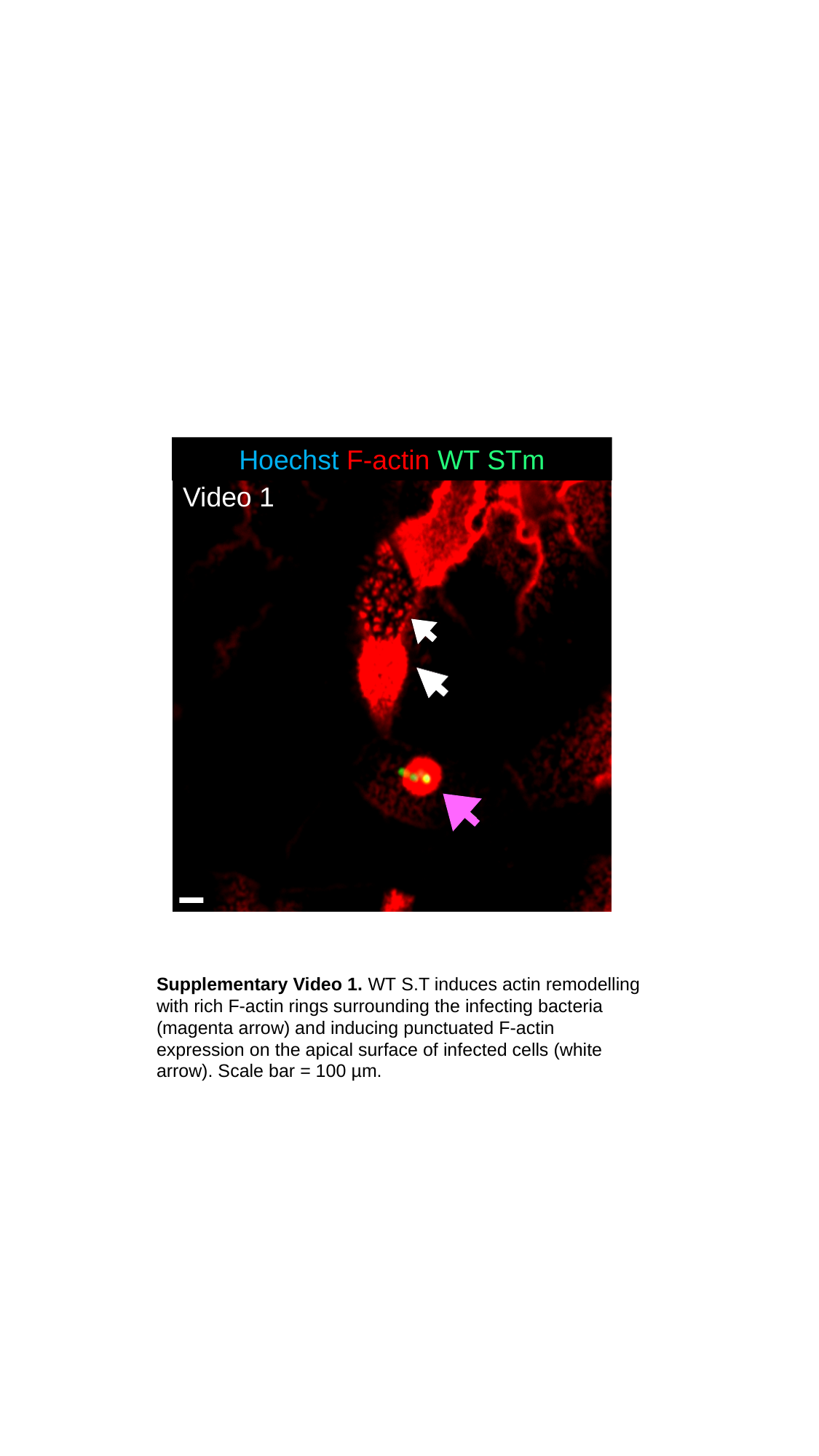

Hoechst F-actin WT STm
Video 1
Video
Supplementary Video 1. WT S.T induces actin remodelling with rich F-actin rings surrounding the infecting bacteria (magenta arrow) and inducing punctuated F-actin expression on the apical surface of infected cells (white arrow). Scale bar = 100 µm.
