## Supplementary Video 2 for "Disentangling the Innate Immune Responses of Intestinal Epithelial Cells and Lamina Propria Cells to *Salmonella* Typhimurium Infection in Chickens"

### Slide 1
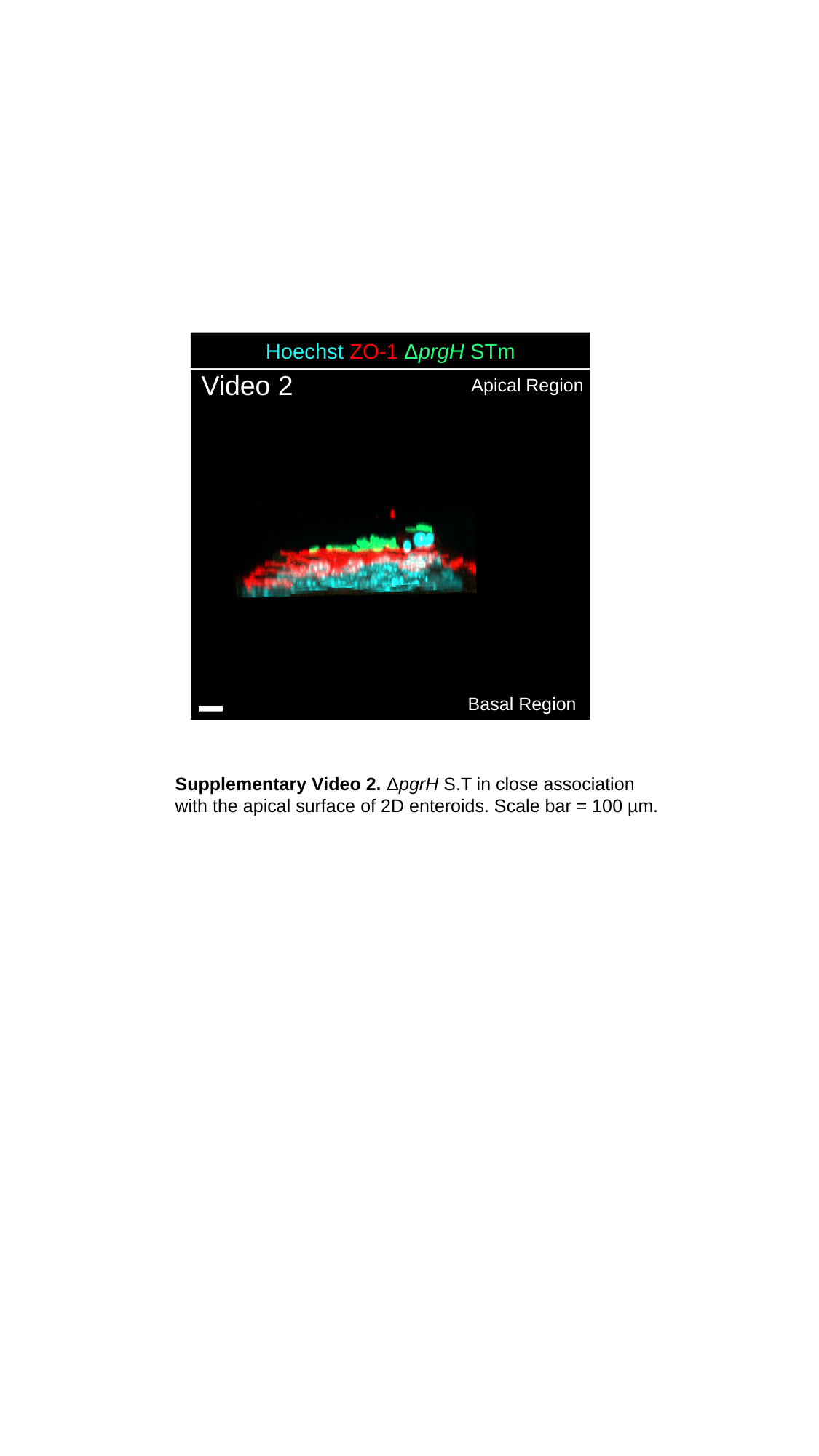

Hoechst ZO-1 ΔprgH STm
Video 2
Apical Region
Basal Region
Supplementary Video 2. ΔpgrH S.T in close association with the apical surface of 2D enteroids. Scale bar = 100 µm.
Apical surface of 3D enteroid
